# Haploid androgenetic development in bovines reveals imbalanced WNT signaling and impaired cell fate differentiation

**DOI:** 10.1101/2023.01.27.525928

**Authors:** Luis M. Aguila, Ricardo P. Nociti, Rafael V. Sampaio, Jacinthe Therrien, Flavio V. Meirelles, Ricardo N. Felmer, Lawrence C. Smith

## Abstract

Haploid embryos have contributed significantly to our understanding of the role of parental genomes in development and can be applied to important biotechnology for human and animal species. However, development to the blastocyst stage is severely hindered in bovine haploid androgenetic embryos (hAE). To further our understanding of such developmental arrest, we performed a comprehensive comparison of the transcriptomic profile of morula-stage embryos, which were validated by qRT-PCR of transcripts associated with differentiation in haploid and biparental embryos. Among numerous disturbances, results showed that pluripotency pathways, especially the wingless-related integration site (WNT) signaling, were particularly unbalanced in hAE. Moreover, transcript levels of *KLF4, NANOG, POU5F1, SOX2, CDX2, CTNNBL1, AXIN2*, and *GSK3B* were noticeably altered in hAE, suggesting disturbance of pluripotency and canonical WNT pathway. To evaluate the role of WNT on hAE competence, we exposed early day-5 morula stage embryos to the *GSK3B* inhibitor CHIR99021. Although no alterations were observed in pluripotency and WNT-related transcripts, exposure to CHIR99021 improved their ability to reach the blastocysts stage, confirming the importance of the WNT pathway in the developmental features of bovine hAE.

**Summary statement:** This study shows the importance of the WNT pathway on bovine haploid androgenetic development by walking through transcriptomics and pluripotency markers associated with cell fate determination during early development.

## Introduction

Landmark experiments that occurred independently by two groups close to four decades ago lay the ground for our current understanding of the essential and complementary contributions of the maternal and paternal genome in mammalian development (Barton et al., 1984; McGrath & Solter, 1984; Surani et al., 1984). These reports showed that while embryos derived from zygotes with two male pronuclei (diploid androgenotes) cannot develop normally, their trophoblast develops well. Conversely, zygotes with two female pronuclei (diploid parthenotes) can develop rather normal embryos with very poor extraembryonic tissue, indicating that the paternal genome is essential for the development of extraembryonic tissue and the maternal genome is particularly important for the development of the embryo itself. These functional differences in developmental genes between parental genomes founded the epigenetic mechanism of genomic imprinting.

Although recent studies indicated that genetic and epigenetic alterations to specific regions controlling the expression of a few key imprinted genes are sufficient to overcome such barriers to allow complete development to term of both bimaternal and bipaternal mice (Kawahara et al., 2007; Kono et al., 2004; Z. Li et al., 2016; Z.-K. Li et al., 2018; Ogawa et al., 2006; Wei et al., 2022), murine genetically unaltered parthenogenetic and androgenetic embryos die by day 10 and 6.5 of gestation, respectively (Barton et al., 1984; Latham et al., 2002; Surani et al., 1984, 1986). Moreover, it is known that already at the preimplantation stages of development, uniparental embryos are affected with regard to cell numbers, morphology and expression profile in laboratory and domestic species models (Aguila et al., 2021; Cui et al., 2011; Gomez et al., 2009; Kure-bayashi et al., 2000; Lagutina et al., 2004; Latham et al., 1994; Loi et al., 1998; Ozil & Huneau, 2001; Thomson & Solter, 1988; Z. Wang et al., 2008). In cattle, reports have described particularly poor development of androgenetic embryos, indicating that a more thorough investigation of the molecular mechanisms controlling the development of androgenetic embryos is required in this species (Aguila et al., 2021; Lagutina et al., 2004; Vichera et al., 2011; S. Wang et al., 2017; H. Zhang et al., 2014). Importantly, uniparental haploid embryos are very efficient models for genome imprinting research and allow studies on the contribution of the paternal and maternal genome to early embryonic development. Moreover, haploid embryos have been used to derive embryonic stem cells and hold great promise for functional genetic studies and animal biotechnology (Bai et al., 2016, 2019; Kokubu & Takeda, 2014; L. Wang & Li, 2019).

Wingless-related integration site (WNT) signaling is a well-known evolutionary and conserved pathway that regulates crucial aspects of cell fate determination and embryonic development (Krivega et al., 2015). In cattle, there are several studies reporting effects of the activation of WNT signaling during the early period of embryonic development (Aparicio et al., 2010; Denicol et al., 2013). For instance, a recent report observed that activation of WNT signaling by the small molecule CHIR99021 increased the levels of *NANOG* and *OCT4* transcripts and *NANOG* positive cells within the inner-cell-mass (ICM), indicating that WNT activation leads to the formation of the ICM and may significantly impact the pluripotency profile and quality of the resulting blastocysts (Warzych et al., 2020).

Thus, the present study aimed to profile global transcriptomic of bovine haploid embryos and investigate on pluripotency features and signalling pathways associated with early developmental failure. In addition, we examined the effect of activating the WNT pathway using the GSK3b inhibitor CHIR99021 and how it impacts on developmental competence of bovine haploid androgenetic embryos. Transcriptomic analysis points to a role of the WNT and pluripotency pathways leading to early differentiation anomalies that can be alleviated by exposure to CHIR99021. The potential significance of these findings is discussed.

## Material and method

### Oocyte collection and in vitro maturation

Bovine ovaries were obtained from a local slaughterhouse and transported to the laboratory in sterile 0.9% NaCl at 25–30°C in a thermos bottle. Cumulus–oocyte complexes (COCs) were aspirated from 5-10 mm antral follicles using a 12-gauge disposable needle. For in vitro maturation (IVM), COCs with several cumulus cell layers were selected, washed, and placed in a maturation medium composed of TCM199 (Invitrogen Life Technologies), 10% fetal bovine serum (FBS; Invitrogen Life Technologies), 0.2 mM pyruvate (Sigma-Aldrich), 50 mg/mL gentamicin (Sigma-Aldrich), 6 μg/mL luteinizing hormone (Sioux Biochemical), 6 μg/mL follicle-stimulating hormone (Bioniche Life Science) and 1 μg/mL estradiol (Sigma-Aldrich). In vitro oocyte maturation was performed for 22-24 h at 38.5°C in a humidified atmosphere at 5% CO_2_.

### Sperm preparation

Straws of sex-sorted semen stored in liquid nitrogen were thawed for 1 min in a water bath at 35.8°C, added to a discontinuous silane-coated silica gradient (45 over 90% BoviPure, Nidacon Laboratories AB), and centrifuged at 600 X g for 5 min. The supernatant containing the cryoprotectant and dead spermatozoa was discarded, and the pellet with viable spermatozoa was re-suspended in 1 mL of modified Tyrode’s lactate (TL) medium and centrifuged at 300 X g for 2 min.

### Production of biparental embryos

In vitro fertilization (IVF): After 20–24 h of IVM, COCs were washed twice in TL medium before being transferred in groups of 5–48 μl droplets under mineral oil. The IVF droplets consisted of modified TL medium supplemented with fatty-acid-free BSA (0.6% w/v), pyruvic acid (0.2 mM), heparin (2 μg/mL), and gentamycin (50 mg/mL). COCs were transferred to IVF droplets 15 min prior to adding the spermatozoa. To stimulate sperm motility, penicillamine (2 mM; Sigma-Aldrich), hypotaurine (1 mM; Sigma-Aldrich) and epinephrine (250 mM; Sigma-Aldrich) were added to each droplet. The selected spermatozoa were counted using a hemocytometer and diluted with IVF medium to obtain a final concentration of 1 × 10^6^ sperm/mL. Finally, 2 μL of the sperm suspension was added to the droplets containing the matured COCs. The fertilization medium was incubated at 38.5°C for 18 h in a humidified atmosphere of 95% air and 5% CO2. Presumptive zygotes were denuded by gentle pipetting.

Intracytoplasmic sperm injection (ICSI): ICSI was performed according to standard protocols (Horiuchi et al., 2002) on the stage of a Nikon Ti-S inverted microscope (Nikon Canada Inc.,) fitted with Narishige micromanipulators (Narishige International) and a Piezo drill system (PMM 150HJ/FU; Prime tech Ltd.). Before ICSI, oocytes were denuded of granulosa cells by gently pipetting in the presence of 1 mg/mL hyaluronidase, selected for the presence of the first polar body and randomly allocated to experimental groups. After ICSI, oocytes were washed at least three times and cultured in modified synthetic oviduct fluid (mSOF) media as previously described (Landry et al., 2016).

### Production of haploid embryos

Bovine haploid androgenetic embryos (hAE) were produced using female-sorted (X-chromosome carrying) as previously reported by our group (Aguila et al., 2021). Bovine haploid parthenogenetic embryos (hPE) were produced according to Valencia et al., (2021). Briefly, chemical oocyte activation was performed between 20 to 24 h after IVM by 5 min exposure to 5 μM ionomicyn (Calbiochem). To obtain haploid parthenotes, ionomycin treatment was followed by incubation in 10 mg/mL cycloheximide (CHX; Sigma-Aldrich) for 5 h. After parthenogenetic activation, presumptive zygotes were washed and allocated to *in vitro* culture droplets.

### *In vitro* culture

For in vitro culture, groups of 10 embryos were placed in droplets (10 μl) of modified serum-free synthetic oviduct fluid (mSOF) with non-essential amino acids, 3 mM EDTA, and 0.4% fatty-acid-free BSA (Sigma-Aldrich) under embryo-tested mineral oil. The embryo culture dishes were incubated at 38.5°C with 6.5% CO_2_, 5% O_2_, and 88.5% N_2_ in 100% humidity. In some treatment groups, GSK3B-inhibition was induced from day 5 onward with 3 μM of CHIR99021 (TOCRIS) (Tribulo et al., 2017). Morulas cultured in 0.001% of DMSO were used as the vehicle control.

### RNA-seq Library preparation and RNA sequencing

Day-6 morula stage embryos from the haploid androgenetic (hAE), parthenogenetic (hPE) and biparental (ICSI) groups were selected (five of each group) and individually analyzed. To obtain the cDNA samples for sequencing, embryos were transferred with the minimal solution possible (<1μL) to microtubes (free DNAase/RNAase) and snap-frozen individually in liquid nitrogen and stored at −80°C until RNA extraction. The SMART-Seq®HT kit (Takara Bio, USA), which uses a poly A tail filter to capture RNA, was used for RNA extraction, amplification, and cDNA production following the manufacturer’s recommendation. RNA quantification was verified by fluorometry (Qubit® ThermoFischer) and RNA quality control was verified using the Agilent Bioanalyzer system. RNA was amplified in 17 PCR cycles and selected for sequencing based on RNA concentration and integrity. Libraries were prepared using the NextEra XT Stranded mRNA Sample Prep kit and quantified by qPCR using the KAPA Library Quantification kit (Illumina; KAPA Biosystems).

Sequencing was performed on Illumina Nova seq (2×100bp) and reads quality was assessed using FASTQC software (http://www.bioinformatics.babraham.ac.uk/projects/fastqc/) (Andrews et al., 2018). The samples underwent quality filtering (average Phred score > 24 and read length >30) and adapter removal using cutadapt implemented in the Trimgalore pipeline (Martin, 2010). After the quality filter, sequencing reads of each sample were aligned using STAR (Dobin et al., 2013) with standard parameters for alignment with the *Bos taurus* genome (Ensembl and NCBI Bos taurus ARS-UCD1.2), and gene count was analyzed using featureCounts (Liao et al., 2014) implemented in the Rsubread package (Liao & Smyth, 2019).

### Differential gene expression and functional enrichment analysis

Genes were considered expressed when they presented more than 4 counts in at least 4 samples. Differential gene expression analysis was performed using the DESeq2 package (Love et al., 2014), considering significance when the adjusted *p* values were less than 0.10 (Benjamini-Hochberg - “BH”) and the absolute value of log2 foldchange was greater than 0.5. Additionally, we considered genes as differentially expressed if they were exclusive, expressed in one group (at least 5 counts in all technical replicates), and not expressed in the other group (zero counts in all technical replicates) within comparison and using the function filterByExpr from edgeR package (Robinson et al., 2010). We estimate the hub genes using CeTF (Biagi et al., 2021) based on RIF— Regulatory Impact Factor and PCIT—Partial Correlation and Information Theory (Reverter et al., 2010; Reverter & Chan, 2008). Gene ontology analysis was performed using clusterProfiler (Yu et al., 2012) and pathways explored using Pathview (Luo & Brouwer, 2013). Data were visualized using R software, in which we primarily observed the classification, intensity, and difference in expression between groups. Exploratory data analysis was performed with principal component analysis using plotpca function from DESeq2 (Love et al., 2014) and ggplot2 package (Wickham, 2009), smearplots built with ggplot2 (Wickham, 2009), heatmaps using Ward.D2 clusterization method from pheatmap package (Kolde, 2019) and upsetplot using ComplexUpset package (Krassowski, 2020; Lex et al., 2014). Moreover, data was cross-validated using human data previously published (Leng, Sun, & Huang, 2019) and submitted to Gene Expression Omnibus (GEO) under the GSE133856 accession code. The data was downloaded using prefetch from the SRA-toolkit and converted to fastq format. Human data was then aligned, and gene count was performed using the GRCh38.106 Human genome reference and processed according to the above-mentioned pipeline. Gene comparison between human and bovine genes was performed using only the human homolog genes, which were obtained with biomart (Durinck et al., 2005, 2011).

### RNA extraction and RT-PCR

For analysis of gene expression, Day-6 morula stage embryos were pooled in groups of 5. Blastocysts were analyzed individually. Each group was carried out in at least three biological replicates and each replicate was run in duplicate. Total RNA was extracted using the Arcturus PicoPure RNA Isolation kit (Life technologies) and reverse transcribed into cDNA using SuperScript Vilo (Invitrogen). Semi-quantitative RT-PCR was performed using the RotorGene SyBr Green PCR kit (Qiagen) in a Rotorgene Q PCR cycler under the following amplification conditions: 95°C for 5 min, followed by 40 cycles at 95°C for 5 secs and at 60°C for 10 secs. Primers were designed using Oligo6 software and the geometric means of three reference genes (*GAPDH, ACTB* and *SF3A*) were used for normalization. The stability of the reference genes across our samples was confirmed using Bestkeeper (Pfaffl et al., 2004). The list of all primers used can be found in **Table S1**.

### Developmental potential and embryo quality evaluation

The developmental potential was assessed as previously reported (Águila et al., 2017). Briefly, cleavage rate was recorded at 48 hours post fertilization (hpf), while morula and blastocyst development were recorded on days 6 and 8 post-fertilization, respectively. After assessment of development, embryos were either fixed for cell number evaluation or snap frozen in liquid N_2_ and stored at −80 °C for RNA extraction. Embryo quality was assessed based on morphology and total cell number. Briefly, embryos at day 8 were classified morphologically as morula (compacted >30 cells), early blastocyst (beginning of a blastocoel cavity), expanded blastocyst (cavity larger than the embryo and zona pellucida thinning), and hatched blastocyst (complete extrusion from zona pellucida).

### Immunostaining

Immunostaining was performed as described previously (Sampaio et al., 2020). Embryos were collected and washed in PBS with PVA and fixed with 4% paraformaldehyde for 15 min and permeabilized with D-PBS with 1% Triton X-100 for 30 min. After blocking for 2 h in D-PBS with 0.1% Triton X-100, 1% BSA, and 5% goat serum (Gibco, NZ), embryos were placed in primary antibody solution consisting of blocking buffer, a mouse antibody anti-Sox2 (Abcam ab10005 at 1:500) and rabbit antibody anti-Cdx2 (Abcam ab10305 at 1:500) overnight at 4 °C. After washing 3 × for 10 min and 3 × for 20 min each, embryos were incubated with secondary antibodies (1:2000) Alexa Fluor 594-conjugated goat anti-rabbit IgG (Life Tech, cat. # A-11036) and Alexa Fluor 488-conjugate goat anti-mouse IgG (Life Tech, cat. #: A-11029) both at RT for 1 h. Finally, embryos were washed 3× for 10 min and 3× for 20 min each, mounted on slides with Prolong Gold Antifade with DAPI (Life Tech, cat. # P36935), and evaluated using confocal microscopy (Olympus FV 1000 laser-scanning confocal microscope).

### Statistical analysis

Quantitative data sets are presented as means and standard deviation (± S.D) and analyzed using one-way ANOVA. Post hoc analysis to identify differences between groups was performed using Tukey test. Binomial data sets, such as pronuclear formation, were analyzed by using Fisher test. Differences were considered significant at *p* < 0.05. Figures and statistical analysis were obtained using the software R (https://www.R-project.org/).

## Results

### Global transcriptomic features of day-6 morula-stage haploid embryos

To examine gene expression patterns of uniparental haploid (androgenetic and parthenogenetic) bovine embryos, we sequenced the transcriptome from individual morula stage embryos, including biparental (BI) ICSI-derived embryos as control. In total, we obtained about 373.9×10^6^ reads of clean data from 5 individual embryos per group (**Fig. S1**). The mapping rate was above 85% using STAR with bovine reference genome ARS-UCD1.2 from ENSEMBL and NCBI (redundant genes were removed to build only one count table). After quality control, a total of **21941 genes** were considered as expressed transcripts. We performed clustering of haploid and biparental (ICSI) embryos using principal component analysis (PCA), heatmapping, and volcano plots. In addition, High Pearson correlation coefficients were found among biological replicates with R-values of 0.954. 0.944 and 0.887 for BI, hPE, and hAE respectively, demonstrating the reproducibility of sample preparation and the sequencing protocols. While PCA revealed clear grouping between hPE and biparental ICSI embryos, hAE samples were clearly unrelated to the samples from the other groups (**Fig. 1A**). Differential expression analysis between the hAE and ICSI samples identified 2347 differentially expressed genes (DEGs), 12 genes were exclusively expressed in ICSI, 432 mRNAs and 18 transcriptional factors (TFs) were considered hub genes between androgenotes and ICSI embryos (**Fig. 1B**). When comparing androgenotes versus parthenotes we identified 1523 DEGs, 1 gene (LOC100849023) that was exclusively expressed in hAE, 6 genes were exclusively expressed in hPE, 447 mRNAs and 17 TFs that were considered hub genes (**Fig. 1C**). Comparisons of parthenotes and biparental groups identified 55 DEGs, of which 5 genes were exclusively expressed, 238 mRNAs and 7 TFs hub genes. In general, the number of varying genes between parthenotes and ICSI was smaller when compared hAE against biparental ICSI or hPE, indicating a higher level of discordance of the androgenetic group. To identify shared regulations, we compared the sets of more expressed genes obtained from each group (**Fig. 1E**). More than 60% (1602/2608) of DEGs were present in androgenotes, 16% (415/2608) in ICSI and only 8% (202/2608) in hPE. The highest intersection was found between transcriptomes of ICSI and hPE (345 genes, representing 45% and 65% of the total number of up-regulated genes for ICSI and parthenotes, respectively) (**Fig. 1E**). In contrast, the intersection between transcriptomes of ICSI and hAE was considerably lower (only 33 genes representing 4% and 2% of the total number of more expressed genes for ICSI and androgenotes, respectively) (**Fig. 1E**). In a similar fashion, the intersection between hPE and hAE was even smaller (11 genes, representing 2% and 0.7% of the total number of more expressed genes for parthenotes and androgenotes, respectively) (**Fig. 1E**). Moreover, hierarchical clustering of DEGs showed a similar clustering by the heat map between the hPE and BI embryos and a clear contrast to the hAE group (**Fig. 1F**). Heatmap of top fifty DEGs between biparental and haploid androgenetic embryos is shown in **Fig. 1G**. Among the top fifty more DEGs between parthenotes and biparental embryos, the maternally expressed imprinted gene *MEG3* was more represented in parthenotes compared to ICSI. In contrast, imprinted genes *SNRPN* and *SNURF*, both expressed exclusively from the paternal allele, were highly expressed in androgenotes and biparental groups compared to hPE. Altogether, a differential transcriptional profile of imprinted genes in hAE and hPE supports their respective parental origin.

**Figure 1.**
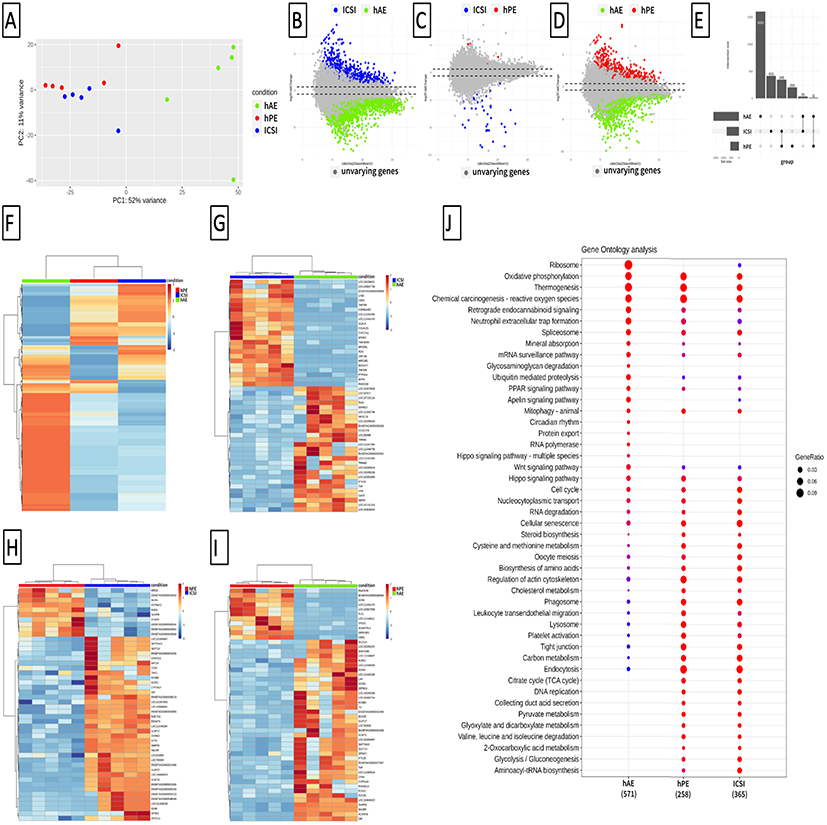
The uniparental haploid genome influences transcriptomic profile at the morula stage. A, Principal component analysis (PCA) plot based on transcriptomic differences among haploid androgenetic (hAE; green dots), haploid parthenogenetic (hPE; red dots) and biparental (ICSI; blue dots) embryos. B-D, Smear plot depicting differences on expression between (B), haploid androgenetic (hAE; green dots) and biparental biparental (ICSI; blue dots) embryos, (C), haploid parthenogenetic (hPE; red dots) and biparental (ICSI; blue dots) embryos, and (D), haploid androgenetic (hAE; green dots) and haploid parthenogenetic (hPE; red dots) embryos; dots in grey are not differentially expressed based in the filterByExpr function from edgeR package. E, Bar chart illustrating the most represented transcripts by each group of embryos and intersection size between transcriptomes. F-I, Heatmap illustration showing top fifty differentially expressed genes among haploid and biparental embryos. J, Gene ontology enrichment analysis of the 1194 more expressed genes in transcriptomic from haploid androgenetic (hAE; green dots), haploid parthenogenetic (hPE; red dots) and biparental (ICSI; blue dots) embryos. The red and blue represent enrichment (absolute value of log2 foldchange greater than 0.5) for up- and downregulated DEGs, respectively.

A total of 1194 DEGS were identified in the KEGG pathways ontology analysis. Comparative KO analysis between the androgenote and control groups revealed enrichment of genes participating in the ribosome, glycosaminoglycan degradation, ubiquitin mediated proteolysis, apeling signalling, protein export, RNA polymerase, and interestingly pathways related to embryonic development (Hippo and WNT signaling). In contrast, haploid androgenetic embryos showed a loss of pathways related to cellular metabolism (steroid biosynthesis, cysteine, and methionine metabolism, biosynthesis of amino acids, cholesterol metabolism, citrate cycle, pyruvate metabolism, among others), regulation of actin cytoskeleton, phagosome, tight junction, and DNA replication activities compared to control groups (**Fig. 1J**). Overall, these results suggest a dramatic influence of the parental genome of haploid morula stage embryos on their transcriptomic profile and reveal that while biparental embryos often share the transcriptomic profile of parthenogenetic embryos, androgenotes show substantial quantitative and qualitative differences in their transcripts during early embryogenesis.

### Unbalanced signaling regulating pluripotency in uniparental haploid embryos

Haploid androgenetic development in the bovine is characterized by poor morphological quality associated with a lower potency to expand and form blastocele (Aguila et al., 2021; Vichera et al., 2011). Thus, we were interested in obtaining deeper insights regarding the pluripotency features of bhAE. Again, the PCA analysis revealed that androgenetic samples presented the most diverse clustering pattern, while both parthenogenetic and biparental samples were similar (**Fig. 2A-C**). In the same fashion, heatmap analysis showed a homogeneous profile between biparental and parthenotes, but a differential for hAE, where the counting of GSK3B, FZD1, ID4, ID2, ESRRB, TCF7, JAK1, and ISL1 transcripts was higher (**Fig. 2D**). Pathview analysis showed that core factors JAK, BMP4 and GSK3B were overexpressed and FGF less active in androgenotes compared to control embryos (**Fig. 2E-F**). Instead, core factors AKT, BMP4, FGF and GSK3B were more expressed in biparental ICSI embryos, but fewer amounts of JAK, BMPR, TCF1, ID and ESRRB were found when compared to parthenogenetic samples (**Fig. 2G**). Interestingly, the well-known TE lineage markers (CDX2 and DAB2) as well as ICM markers (POU5F1 and NANOG) were not among the most variable genes, indicating nascent TE and ICM lineages. These data suggest unbalanced pluripotency of uniparental haploid embryos, a potential link to their inefficient ability to undergo blastulation.

**Figure 2.**
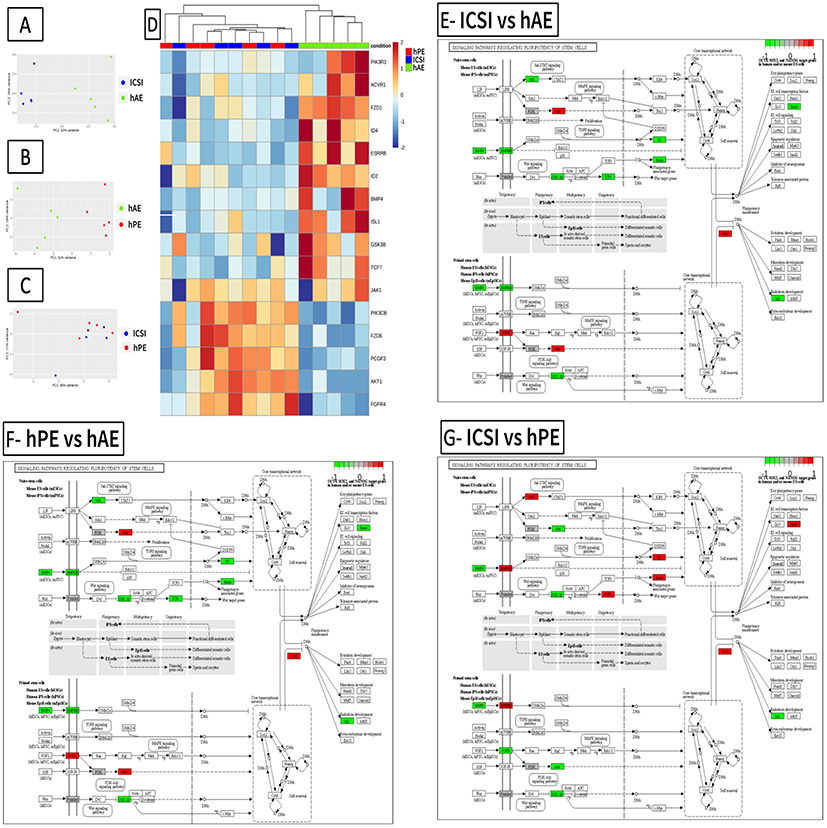
Haploid androgenetic embryos exhibit differential levels of pluripotency factors at the morula stage. A-C, Principal component analysis (PCA) plot based on transcriptomic profiles of pluripotency-related factors between (A), haploid androgenetic (hAE; green dots) and biparental (ICSI; blue dots) embryos, (B), haploid parthenogenetic (hPE; red dots) and haploid androgenetic (hAE; green dots) embryos, and (D), haploid parthenogenetic (hPE; red dots) and biparental (ICSI; blue dots) embryos. D, Heatmap showing the expression levels of factors associated with pathways regulating pluripotency of stem cells. Using hierarchical clustering, genes are segregated into two groups, where haploid parthenogenetic (hPE) samples grouped with biparental (ICSI) embryos, distinctly than haploid androgenetic (hAE) counterparts. E-G, differential expression levels of signaling factors regulating pluripotency networks between biparental (ICSI) and haploid androgenetic (hAE) embryos (E), haploid parthenogenetic (hPE) and haploid androgenetic (hAE) embryos (F), and biparental (ICSI) and haploid parthenogenetic (hPE) embryos (F).

The WNT (Wingless-related integration site) pathway regulates crucial aspects of cell fate determination and embryonic development. Previous studies have shown that WNT activation via blocking GSK3B from the morula stage onwards improves blastocyst morphology and epiblast-specific gene expression (Harris et al 2013; Madeja et al 2015). To further analyze whether the developmental constraints of uniparental haploid embryos are associated to altered pluripotency, we focused our analytical pipeline on the expression profiles of WNT-related genes. The PCA was segregated differentially in androgenetic samples compared to biparetal and parthenogenetic groups (**Figure 3 A-C**). Moreover, the heatmap evidenced a strong expression of WNT genes in androgenotes, particularly for GSK3B, suggesting an alteration to the WNT pathway (**Fig. 3D**). Once again, path-view indicates a differential expression of key factors of the canonical (RSPO, FRP, GSK3B, TCF/LEF, gamma-Catenin, and PPAR gamma), p53 signaling (Siah1), planar cell polarity (Daam and RhoA) and calcium-dependent (NFAT) Wnt pathways in hAE compared to ICSI and parthenotes (**Fig. 3E, F**). Finally, pathway analysis also showed overrepresentation of canonical (BAMBI, GSK3B) and planar cell polarity (Daam1 and RhoA) signalling in biparental ICSI compared to hPE (**Fig. 3G)**. Together, the RNAseq analysis indicates higher heterogeneity in WNT activity among groups, predominantly for GSK3B.

**Figure 3.**
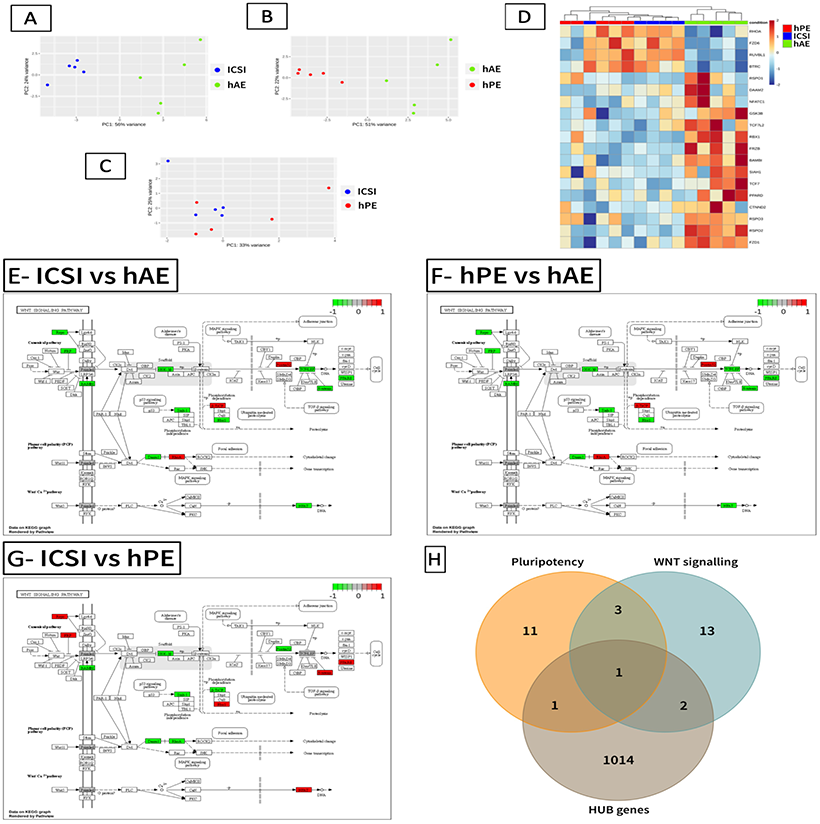
Haploid androgenetic embryos display differential expression of WNT signaling factors at the morula stage. A-C, Principal component analysis (PCA) plot based on transcriptomic profiles of WNT-signaling factors between (A), haploid androgenetic (green dots) and biparental ICSI (blue dots) embryos, (B), haploid parthenogenetic (red dots) and haploid androgenetic (green dots) embryos, and (C), haploid parthenogenetic (red dots) and biparental ICSI (blue dots) embryos. D, Heatmap showing the relative expression levels of factors regulating WNT-signaling. Using hierarchical clustering, genes are segregated into two groups, where haploid parthenogenetic (hPE) samples grouped with biparental (ICSI) embryos, distinctly than haploid androgenetic (hAE) counterparts. E-G, differential expression levels of signaling factors regulating WNT network pathway between biparental (ICSI) and haploid androgenetic (hAE) embryos (E), haploid parthenogenetic (hPE) and haploid androgenetic (hAE) embryos (F), and biparental (ICSI) and haploid parthenogenetic (hPE) embryos (F). H, Venn diagram illustrating relationship between pluripotency, WNT signalling and hub genes of bovine haploid androgenetic morula-stage embryos.

To identify genes that are central and highly connected to pluripotency networks, we conducted hub gene identification analysis using the CeTF package. We assigned all protein coding genes (mRNAs) with an absolute RIF value > 2 as hub genes. These analyses identified 4 genes from the WNT-pathway associated with pluripotency, 3 genes from the WNT-pathway related to Hub genes, and 2 genes associated with pluripotency were related to Hub genes. Importantly, the only gene from the WNT-signalling pathway that was associated with pluripotency and Hub genes was GSK3B (**Fig. 3H**). Together, these data indicated unbalanced pluripotency signaling, but also identified potential key regulators of cell differentiation, highlighting those transcriptomic differences, could be associated with the poor developmental competence observed in haploid phenotypes.

### Uniparental human and bovine morula-stage embryos share analogous imprinting profile

Next, to corroborate our finding, we performed in silico validation via bioinformatic analysis of previously published transcriptomic data of human uniparental (diploid) and biparental (ICSI) morula-stage embryos (Leng, Sun, Huang, et al., 2019). PCA revealed a similar clustering pattern between hPE and biparental embryos, and again androgenetic samples were grouped more distant from the other groups (Fig. 4A). The heatmap of human homologous genes revealed a closer profile between biparental and diploid androgenetic embryos, while diploid parthenogenetic embryos differed from the other groups (**Fig. 4B, C)**. In contrast, our bovine data indicated a clear similarity between biparental ICSI and hPE, while hAE displayed a different profile (**Fig. 4B, C**). Due to the dissimilar transcriptomic pattern between bovine and human samples, we focused our analysis on the expression of imprinted genes. Uniparental embryos possess only one (haploid) or two (diploid) copies of either the paternal or maternal genomes and, therefore, lack gene transcripts expressed monoallelically from either one or the other. This analysis showed analogous clustering of imprinted genes in bovine and human data. For instance, the paternally expressed SNRPN, PEG10, PLAGL1, and KCNQ1OT1 genes were present abundantly in androgenetic but barely expressed in parthenogenetic samples (**Fig. 4D, E**). On the opposite, maternally expressed genes such as MEG8, MEG3, and GAB1 were more present in parthenogenetic rather than androgenetic samples (**Fig. 4D, E**). Thus, these data indicate that uniparental human and bovine embryos share similar imprinting profiles, regardless of their ploidy condition.

**Figure 4.**
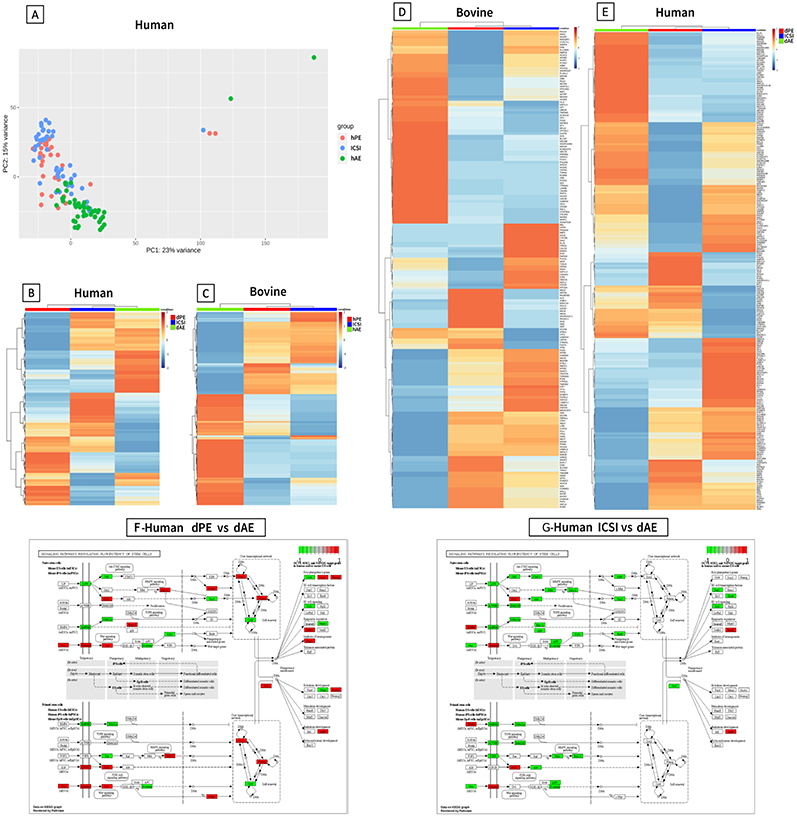
The transcriptomic profile of human uniparental diploid morula-stage embryos behaves differentially regarding to bovine species but share global imprinted patterns. B-C, Heatmap illustration showing global transcriptomic profiles of homologous genes from (B) human diploid parthenogenetic (dPE), diploid androgenetic (dAE) and biparental (ICSI) embryos, and (C) bovine haploid parthenogenetic (hPE), haploid androgenetic (hAE) and biparental (ICSI) embryos. D-E, Heatmap illustration showing imprinted transcriptomic profiles of (D) human diploid parthenogenetic (dPE), diploid androgenetic (dAE) and biparental (ICSI) embryos, and (E) bovine haploid parthenogenetic (hPE), haploid androgenetic (hAE) and biparental (ICSI) embryos. F-G, signaling factors regulating pluripotency networks between human diploid parthenogenetic (dPE) and diploid androgenetic (dAE) embryos (F), and between biparental (ICSI) and diploid androgenetic (dAE) (G).

We next sought to examine signaling pathways regulating pluripotency in uniparental human data by applying our bioinformatics pipeline. Contrary to bovine data, we found that PIK3, ERK1/2, and Dvl pathways were more active in human androgenetic samples than in diploid parthenotes. Noticeably, GSK3B was not active, and B-catenin was less represented in diploid androgenotes (**Fig. 4F**). On the other hand, diploid androgenetic data showed that most of the signaling pathways were less active (JAK/STAT3, SMADs, Wnt/B-catenin, Mek/ERK) when compared to ICSI (**Fig. 4G**). Altogether, bioinformatic *in silico* analysis of human published data indicated that uniparental androgenetic diploid embryos display different transcriptomic patterns compared to bovine hAE, while the imprinting profile is relatively conserved between species.

### Pluripotency and WNT-associated transcripts are altered in haploid androgenetic day-6 morula stage embryos

To further validate results obtained by transcriptomic analysis, we analyzed the expression of pluripotency and WNT-related genes via qRT-PCR. We showed that NANOG, KFL4, and GSK3B were upregulated, but also CDX2 and AXIN2 were downregulated in hAE morula embryos when compared to hPE and biparental ICSI (**Fig. 5A**). SOX2 and CTNNBL1 showed overexpression in hAE but only when compared to biparental groups (**Fig. 5A**). Interestingly, ICSI and hPE that remained at the morula stage at day-7 did not show major differences with androgenotes, indicating a resemblance among developmentally retarded hAE and biparental and hPE groups (**Fig. 5B**). These results confirm that morula-stage day-6 hAE have unbalanced pluripotency and a dysregulated expression of the WNT-pathway factors CTNNBL1, GSK3B and AXIN2.

**Figure 5.**
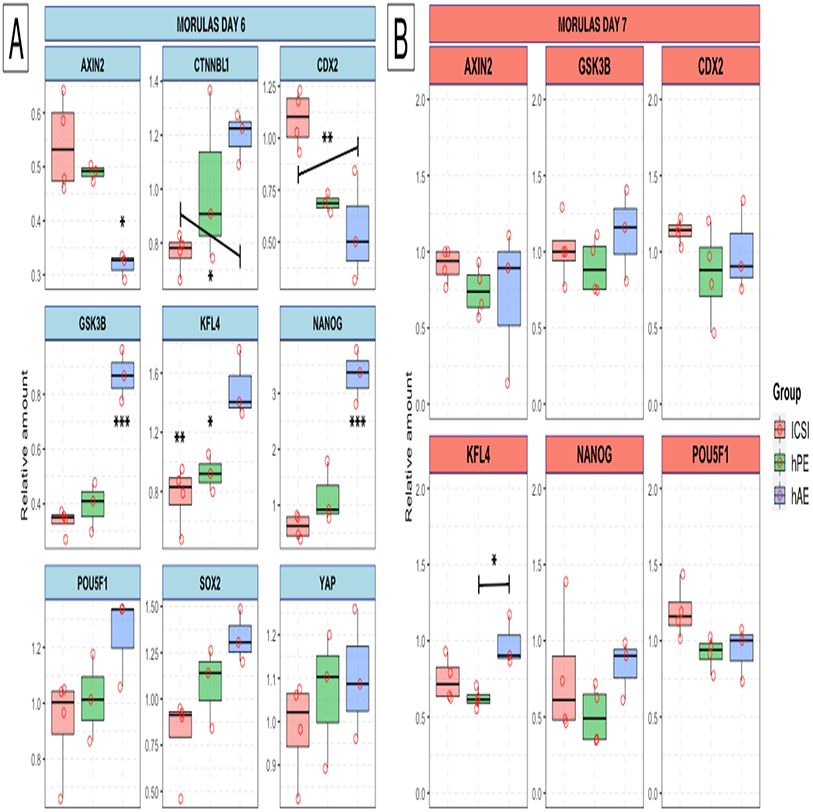
Relative transcript levels of genes associated with pluripotency from biparental (salmon square), haploid parthenogenetic (green square), and haploid androgenetic (light-blue square) at the morula-stage. A, Light-blue boxes: morulas at day 6 post fertilization; B, coral boxes: “arrested” morulas at day 7 post fertilization. Axis inhibition protein 2 (*AXIN2*); Beta-catenin-like protein 1 (*CTNNBL1*); caudal type homeobox 2 (*CDX2*); Glycogen synthase kinase-3 beta (GSK3B); kruppel-like factor 4 (KFL4,); homeobox protein NANOG (*NANOG*); POU domain, class 5, transcription factor 1 (*POU5F1*); sex determining region Y-box 2 (*SOX2*); yes-associated protein 1 (*YAP*). (*p < 0.05, **p < 0.01, ***p < 0.001).

### GSK3B inhibition improves the development in vitro of haploid androgenetic embryos

WNT signalling promotes the expression of key pluripotency-related genes and the stabilization of ICM lineage in bovine embryos (Madeja et al., 2015). Because the transcript analysis revealed the overexpression of GSK3B in bhAE concurrent with a failure to form competent morula at day-6, we evaluated the effect of exposing haploid androgenetic embryos from day 5 onward to the GSK3B-inhibitor CHIR99021 on their developmental potential. Apart from the unexposed control groups, a DMSO control was used as a vehicle control. In the absence of CHIR99021, morula and blastocyst development were lower in the hAE compared to biparental and hPE (**Table 1**). However, when hAE morula-stage embryos were cultured in presence of CHIR99021, the proportion of blastocyst increased significantly compared to the vehicle-control group (78% vs 31% for CHIR99021 and DMSO, respectively), enabling a similar blastulation rate (79%) to control groups (78% and 75% for ICSI and hPE, respectively) (**Table 1)**. Additionally, although morphology was not affected by GSK3B inhibition in biparental and haploid parthenogenetic embryos (**Table S2**), blastocyst morphology of hAE was remarkably improved as indicated by the presence of expanding/expanded embryos that were not observed in the absence of CHIR99021 (**Fig. 6 A,B**, **Table S2**). Thus, inhibition of GSK3B from the morula stage onward enhances the developmental competence of bovine hAE.

**Table 1.** In vitro development of biparental, haploid parthenogenetic and haploid androgenetic embryos cultured in the presence of the GSK3B inhibitor CHIR99021 or without inhibitor (DMSO).

| Group | Oocytes | (n) | Cleaved embryos (%) | Morulas (%) | Blastocyst (%) | Blastocyst/morulas (%) |
| --- | --- | --- | --- | --- | --- | --- |
| **Biparental (ICSI)** | **450** | **(15)** | **270 (60%)** | **106 (24%)<sup>a</sup>** | **84 (19%)<sup>a</sup>** | **(79%)<sup>a</sup>** |
| + DMSO |  |  |  | 82 n.a | 64 n.a | (78%)<sup>a</sup> |
| + Chir99021 |  |  |  | 24 n.a | 20 n.a | (83%)<sup>a</sup> |
| **hPE** | **672** | **(17)** | **420 (63%)** | **114 (17%)<sup>b</sup>** | **86 (12%)<sup>b</sup>** | **(75%)<sup>a</sup>** |
| + DMSO |  |  |  | 83 n.a | 62 n.a | (75%)<sup>a</sup> |
| + Chir99021 |  |  |  | 31 n.a | 24 n.a | (77%)<sup>a</sup> |
| **hAE** | **888** | **(20)** | **603 (68%)** | **79 (9%)<sup>c</sup>** | **42 (5%)<sup>c</sup>** | **(53%)<sup>ab</sup>** |
| + DMSO |  |  |  | 42 n.a | 13 n.a | (31%)<sup>b</sup> |
| + Chir99021 |  |  |  | 37 n.a | 29 n.a | (78%)<sup>a</sup> |

Development to cleavage and blastocyst stages at Day-2 (48 h) and Day-8 (192 h) post fertilization, respectively. ICSI, intracytoplasmic sperm injection. hPE, haploid parthenogenetic embryo obtained by oocyte activation using ionomycin followed by cycloheximide. hAE, haploid androgenetic embryos were obtained by ICSI followed by TII enucleation. n.a.: cleaving and morula's rates were not included because DMSO and CHIR99021 treatment initiated at day 5 of culture.

**Figure 6.**
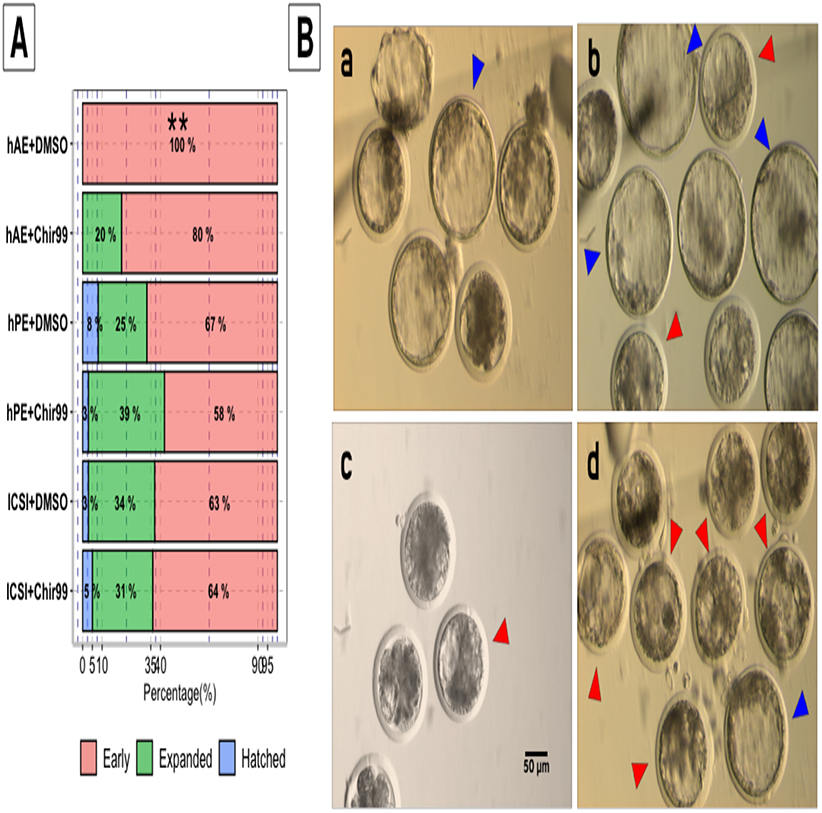
Blastocyst morphology at 192 h of *in vitro* culture. A, percentages of biparental (ICSI+DMSO and ICSI+Chir99), haploid parthenogenetic (hPE+DMSO and hPE+Chir99) and haploid androgenetic (hAE+DMSO and hAE+Chir99) development cultured in the absence (DMSO vehicle) or presence of CHIR99021. B, representative images of a) biparental (ICSI), b) haploid parthenogenetic (hPE), c) haploid androgenetic+DMSO (hAE+DMSO), and d) haploid androgenetic + CHIR99021 (hAE+Chir99) embryos at Day-8 (192 h) of culture. ICSI, intracytoplasmic sperm injection; hPE, haploid parthenogenetic embryo; hAE, haploid androgenetic embryo. Red arrowheads indicate early blastocyst stage. Blue arrowheads indicate expanded/expanding blastocyst stage.

### Transcripts of pluripotency and WNT-related genes are unaffected by GSK3B inhibition

To investigate the mechanisms involved in the improvement of haploid androgenetic development caused by GSK3B inhibition, we compared the transcript levels of pluripotency and WNT-related genes in day-8 blastocyst stage embryos cultured in CHIR99021 (GSK3B-inhibition) via RT-PCR. First, we corroborate that DMSO (0.001% v/v) and CHIR99021 did not affect expression of reference genes used for normalizations (**Fig. S1**). We next analyzed the expression of the same panel of pluripotency-related genes evaluated at the morula stage, but including blastocysts produced by IVF (females) as control. Compared to biparental and parthenotes, GSK3B expression was higher (*p*<0.05) in hAE, both in the presence and absence of CHIR99021 (**Fig. 7**), indicating that the inhibition of GSK3B does not interfere with the transcriptional levels of the gene. Apart from POU5F1, no differences were observed between hAE and biparental embryos (**Fig. 7)**. Although NANOG transcript were not affected by CHIR99021 in any group, a variable overexpression was observed among haploid androgenetic in comparison to parthenogenetic embryos when cultured in presence of DMSO (**Fig. 7)**. Overall, these results indicate that the developmental improvement resulting from the exposure of hAE to GSK3B inhibition does not rely on changes in the levels of key factors involved in embryonic pluripotency and/or WNT-signaling pathway.

**Figure 7.**
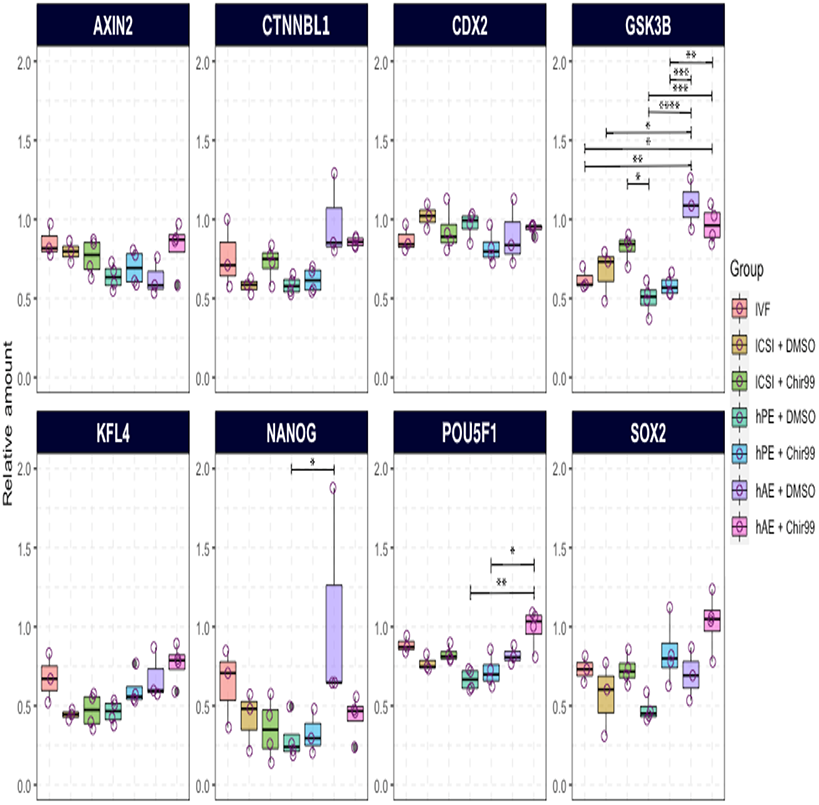
Relative levels of transcripts associated with pluripotency from blastocyst stage embryos (day 8) obtained by *in vitro* fertilization (IVF), intracytoplasmic sperm injection (ICSI), haploid parthenogenetic activation (hPE) or haploid androgenetic (hAE) development, cultured in absence (DMSO) or presence of Chir99021 (Chir99). Salmon squares: IVF; mustard squares: ICSI+DMSO, ICSI embryos cultured in presence of 0.001% DMSO; olive-green squares: ICSI+Chir99, ICSI embryos cultured in presence of Chir99; teal squares: hPE + DMSO, haploid parthenogenetic embryos cultured in presence of 0.001% DMSO; light-blue squares: hPE + Chir99, haploid parthenogenetic embryos cultured in presence of Chir99; purple squares: hAE + DMSO, haploid androgenetic embryos cultured in presence of 0.001% DMSO; magenta squares: hAE + Chir99, haploid androgenetic embryos cultured in presence of Chir99021. Axis inhibition protein 2 (*AXIN2*); Beta-catenin-like protein 1 (*CTNNBL1*); caudal type homeobox 2 (*CDX2*); Glycogen synthase kinase-3 beta (GSK3B); kruppel-like factor 4 (KFL4,); homeobox protein NANOG (*NANOG*); POU domain, class 5, transcription factor 1 (*POU5F1*); sex determining region Y-box 2 (*SOX2*); yes-associated protein 1 (*YAP*). (*p < 0.05, **p < 0.01, ***p < 0.001).

### Low ICM:TE ratio of hAE cannot be rescued by GSK3B-inhibition

To further unravel the role of GSK3B on the developmental competence of haploid androgenotes, we compared total cell number and allocation to inner-cell-mass (ICM) and trophectoderm (TE) in hAE blastocyst exposed or not to CHIR99021. To do so, we performed immunostaining to SOX2 and CDX2, known bovine specific markers of the ICM and TE cells, respectively. Biparental (ICSI) and hPE were used as controls. Regardless of the presence of CHIR99021, hAE contained fewer total cell numbers (DNA-stain), and fewer SOX2- and CDX-2 positive cells (**Fig. 8A, B**). However, the proportional representation of ICM:TE ratio was significantly lower in androgenetic embryos in relation to total cells when compared to ICSI and parthenogenetic blastocysts (**Fig. 8A, B**). On the other hand, CDX2-positive cells were present in higher percentages (around 90%) in hAE than in hPE or ICSI (**Fig. 8A, B**). Moreover, no difference in the amount or proportion of SOX2-expressing cells at the blastocyst stage exists between hAE exposed or not to CHIR99021 (**Fig. 8B**). The limited presence of SOX2-positive cells in androgenetic development was further evidenced on day 10 hAE cultured in the presence of GSK3B inhibitor CHIR99021 (**Fig. 8B**). Thus, these results show that hAE privilege TE differentiation during blastulation and that GSK3B inhibition cannot rescue the embryos to form a proper ICM:TE ratio.

**Figure 8.**
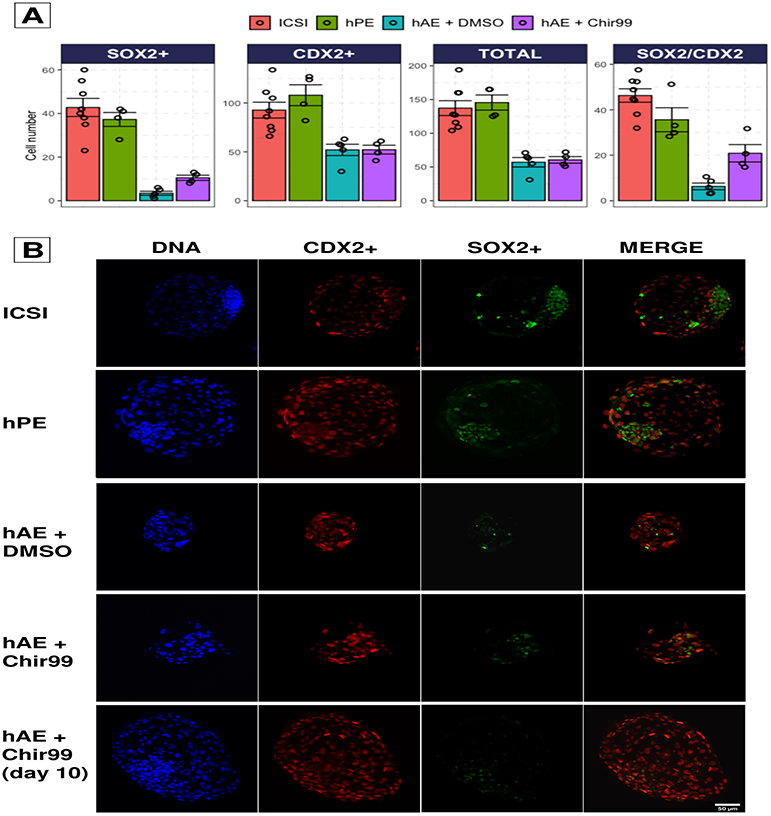
Cell number and allocation of biparental and haploid blastocyst stage embryos. A, Nuclear counts of blastocyst stage embryos harvested at day-8 of culture. B, Representative fluorescent images of blastocyst harvested at day-8 or day-10 of culture. ICSI, intracytoplasmic sperm injection; hPE, haploid parthenogenetic embryo; hAE + DMSO, haploid androgenetic embryos cultured in presence of 0.001% DMSO; hAE + Chir99, haploid androgenetic embryos cultured in presence of Chir99021.

## Discussion

We hereby have performed a comprehensive analysis of the transcriptomic profiles of uniparental haploid and biparental bovine embryos. Our data showed that biparental and hPE embryos share a closer transcriptomic profile at the morula stage when compared to haploid androgenetic transcriptome. Besides, the main pathways associated with pluripotency are unbalanced in haploid androgenetic samples, particularly for genes associated with WNT signaling. Moreover, we show that GSK3B-inhibition enhances the developmental potential of haploid androgenetic morula-stage embryos. Finally, haploid androgenetic blastocysts have lower quality in terms of cell number and ICM formation, indicating a preferential differentiation towards TE lineage that cannot be reversed by GSK3B-inhibition.

Previous studies in mammals have established that the androgenetic embryos have reduced developmental competence from the morula to the blastocyst stage (Aguila et al., 2021; Hu et al., 2015; Lagutina et al., 2004; Latham et al., 2002; Matsukawa et al., 2007; Vichera et al., 2011; S. Wang et al., 2017; Xiao et al., 2013; H. Zhang et al., 2014). Moreover, androgenetic embryos that progress beyond ZGA underwent a second developmental arrest at the morula stage (Aguila et al., 2021). Here, we hypothesized that developmental restriction of bovine morula-stage hAE is associated with poor pluripotency. We first investigated the global transcriptomic profile in uniparental haploid and biparental samples. As reported for human uniparental embryos (Leng, Sun, Huang, et al., 2019), our analysis showed that biparental and parthenogenetic embryos share similar transcriptomic profiles during early embryonic development. Although it is possible that some oocyte-derived transcripts were never degraded, androgenetic samples showed highly heterogeneous transcriptomic profiles compared to the ICSI and parthenogenetic groups, indicating that most oocyte-derived RNAs were no longer present at the morula stage. Because the developmental potential of parthenotes is relatively similar to those of sperm fertilized counterparts in several animal models (Cai et al., 2020; Grupen et al., 1999; Mitalipov et al., 2001), including humans (Mai et al., 2007), this finding rises the possibility that once ZGA has occurred, newly-derived transcripts of maternal origin (nascent from maternal alleles) may play a more essential role during early stages of embryo development in mammalian species.

On the other hand, as previously reported (Aguila et al., 2021; Hu et al., 2015; Latham et al., 1994; Ogawa et al., 2006; Sotomaru et al., 2001), we report perturbed imprinting expression patterns during uniparental development with the overexpression of maternally expressed imprinted genes in parthenotes and overexpression of paternally expressed imprinted genes in androgenotes, consistent with the notion that the absence of the reciprocal parental allele leads to the overexpression of imprinted genes. This differential imprinting expression pattern of haploid embryos determines not only the parental origin of the embryo (Daughtry & Mitalipov, 2014; Sritanaudomchai et al., 2010) but also its association with developmental outcomes (Aguila et al., 2021; Bos-Mikich et al., 2016; Latham et al., 2002). For instance, embryos with only the maternal genome show altered expression patterns of key enzymes required for epigenetic reprogramming (Aguila et al., 2021; Kono, 2006; Peng et al., 2015; Sagi et al., 2019)

In a similar fashion, the bioinformatic analysis revealed a differential profile of pluripotency factors, particularly an unbalanced WNT pathway of overrepresented canonical factors. In the bovine species, modulation of the activity of WNT signaling is necessary for development of ESC (Bogliotti et al., 2018; Xiao, Sosa, et al., 2021). In addition, we identified that GSK3B, a key factor of the canonical WNT signalling pathway, is a central hub gene for haploid androgenetic development and may function as a “master regulator” of gene expression and developmental transition during the first cell-fate differentiation (Denicol et al., 2013; Madeja et al., 2015; Tribulo et al., 2017; Warzych et al., 2020).

Nonetheless, our “in silica” cross-species analysis revealed different transcriptomic profiles of homologous genes between human uniparental diploid and bovine haploid embryos at the morula stage. However, imprinting expression appears conserved in uniparental samples across both species. Comparing uniparental and bi-parental embryos by genome-wide technologies is useful for identifying imprinted genes (Sagi et al., 2019; Stelzer et al., 2015). Our results confirmed the imprinting behavior of most of the genes previously described for bovine species, but also raised the existence of potential imprinting loci not described for the bovines but reported as imprinted in humans. Therefore, the expression profiles established in this report can therefore serve as a reference base for bovine species, since the identification of imprinted genes in livestock species lags behind human and mouse data (O’Doherty et al., 2015).

Further investigation into transcripts of a panel of pluripotency ICM/TE specific lineage markers revealed abnormal expression of the pluripotency factor KLF4 and other markers of ICM, (i.e. NANOG, SOX2, and/or the TE marker CDX2), and genes associated with WNT pathway, i.e. GSK3B, AXIN2, and CTNNBL1. The KLF gene family is likely involved in directing gene reprogramming during EGA in bovine embryos (Bogliotti et al., 2020), and KLF4 is specifically required for both ES cell self-renewal and maintenance of pluripotency by regulating NANOG expression (P. Zhang et al., 2010). On the other hand, NANOG, POU5F1, and SOX2 are the core pluripotency transcription factors supporting stem self-renewal and blastocyst potency (Avilion et al., 2003; Bogliotti et al., 2018; Chambers et al., 2003; Mitsui et al., 2003; Sakurai et al., 2016). For instance, disruption of POU5F1 prevented blastocyst formation and was associated with the bovine embryonic arrest at the morula stage (Daigneault et al., 2018). Similarly, the downregulation of SOX2 compromised the expression of NANOG and preimplantation development (Goissis & Cibelli, 2014), and the deletion of NANOG impaired epiblast formation in the bovine ICM (Mitsui et al., 2003; Ortega et al., 2020). Moreover, CDX2 knockdown in bovine blastocysts resulted in poorly elongated embryos due to reduced TE cell proliferation (Berg et al., 2011). Others have indicated that non-competent embryos have an unbalanced overexpression of NANOG and SOX2 (Velásquez et al., 2019). Similarly, bovine embryos with a decreased developmental competence show increased transcription rates of pluripotency markers (Khan et al., 2012). Aberrant POU5F1 and SOX2 expression in bovine cloned blastocysts have also been related to low developmental competence (Hall et al., 2005; Rodríguez-Alvarez et al., 2010, 2013). In this study, we also recorded more similar expressions of pluripotency genes among “less competent” embryos that were arrested at the morula stage. Previous reports and our findings suggest that a balanced expression of pluripotency genes is required during early development, most likely to maintain appropriate regulation of differentiation and cell proliferation. Indeed, the absence of the maternal genome in androgenetic embryos raises the likelihood of unbalanced gene expression not only of imprinted but also of non-imprinted genes.

In mice (Ogawa et al., 2006) and humans (Sagi et al., 2019) androgenetic blastocysts have a lower number of cells and are associated with hindered blastulation. Our previous study also reported a poor blastulation rate (26%) for hAE compared to biparental (80%) or hPE (68%) (Aguila et al., 2021). Notably here, GSK3B inhibition enhanced embryonic competence by increasing the “blastulation rate”, seen as the proportion of morulas becoming blastocysts. This effect may be associated with the inhibition of an unbalanced GSK3B. However, further studies are needed to determine not only the mRNA levels but also the protein levels of GKS3B in hAE samples. These findings agree with others that also report positive effects of GSK3B inhibition on the developmental competence and quality of bovine embryos (Aparicio et al., 2010; Harris et al., 2013; Meng et al., 2015).

Evaluation of the same panel of transcripts at the blastocyst stage indicated subtle differences among hAE (cultured with DMSO or CHIR99021) and control groups. For instance, although GSK3B remained overexpressed in hAE, the other pluripotency-related transcripts remained similar among groups. According to the study of Madeja et. al., (2015), GSK3B inhibition is enough to elevate the expression of POU5F1 and NANOG in both the ICM and TE. In line with these findings, others have indicated that GSK3B inhibition leads to the formation and stabilization of the ICM by promoting the expression of ICM lineage-specific markers POU5F1 and NANOG (Warzych et al., 2020). However, it has been also reported that WNT activation from the morula stage onwards does not have major effects on the ICM compartment (Kuijk et al., 2012). Although we did not observe huge differences at the transcript level, a phenotypic change towards higher morphological quality was recorded under WNT-activated conditions. Therefore, we postulate that bovine hAE are unable to undergo cell fate differentiation during the transition from morula to blastocyst due, at least in part, due to an unbalanced WNT signaling. Our hypothesis is that activation of WNT/b-catenin signaling facilitates the formation of self-renewing pluripotent cell lines from bovine biparental blastocysts (Ozawa et al., 2012). Nonetheless, others have indicated that WNT-activation impairs blastocyst formation and embryo quality (Denicol et al., 2013; Tribulo et al., 2017; Xiao, Amaral, et al., 2021). Moreover, inhibition of WNT signaling was necessary to derive stable pluripotent ESC expressing pluripotency factors SOX2 and POU5F1 (Bogliotti et al., 2018). Nonetheless, it is noteworthy that haploid androgenetic embryos may respond differently to signaling inhibition when compared to biparental embryos since the lack of the maternal genome creates unbalanced signaling pathways during early development. Thus, the precise underlying mechanisms responsible for the actions of WNT activation on the early development of bovine hAE require further elucidation.

Additionally, our immunoassay confirmed that the number of cells expressing SOX2 regarding total and/or CDX2-positive cells were significantly lower in hAE blastocysts. Although GSK3B inhibition was unable to reverse such an anomaly, CHIR99021 exposure appears to shift androgenotes towards an improved ICM:TE ratio. In mice, the absence of SOX2 promotes stem cell specification toward TE lineage (Masui et al., 2007; Tremble et al., 2021). In this line, it is widely known that androgenetic blastomeres preferentially differentiate in trophectoderm (Barton et al., 1984; McGrath & Solter, 1984; Surani et al., 1984). Moreover, WNT/b-catenin signaling can also stimulate trophectoderm differentiation during early embryo development (Denicol et al., 2013; Krivega et al., 2015; Soto et al., 2021; Xiao, Amaral, et al., 2021). In agreement, WNT-YAP/TAZ signaling regulates the differentiation of trophoblast stem cells with properties that resemble the trophectoderm of bovine blastocysts (C. Wang et al., 2019). In addition, a recent report indicated that culturing with CHIR99201 affects the number of SOX2-positive cells in bovine blastocysts (Xiao, Sosa, et al., 2021). Thus, it will be important in future studies to seek strategies capable of improving ICM-specification of bovine hAE. In the same context, the lack of SOX2 protein detected in hAE did not coincide with its mRNA levels. Since the abundance of proteins cannot be accurately predicted from mRNA profiles and changes in mRNA levels can explain at most 40% of the variability in protein levels (Schwanhäusser et al., 2013; Tian et al., 2004), it is not surprising to find that protein abundance of SOX2 did not match its transcripts levels. In fact, studies in mice (Lu et al., 2009) and bovine (Warzych et al., 2020) ESC have also disclosed that overall changes in protein levels are not accompanied by changes in the expression of the analogous mRNAs.

In conclusion, we have shown that the parental genome of haploid embryos severely influences the transcriptomic profile. Furthermore, it is revealed, for the first time, that early biparental development shares a transcriptomic profile closer to parthenogenetic development. In addition, the poor developmental potential and deficient blastulation rate of bovine haploid androgenetic embryos are associated with unbalanced pluripotency and expression of genes associated with the WNT pathway and cell fate differentiation. We have also shown that in cattle, similarly to other mammalian models, androgenetic-derived embryos preferentially differentiate towards the trophectoderm linage, which could be associated with the lack of a defined ICM. Future studies will aim to analyze the effects of activation/inhibition of other signaling pathways involved in cell fate differentiation and pluripotency on the developmental potential of bovine haploid embryos.

## Author Contributions

LA, RPN, JT and LS contributed to the conception and design of the study. LA, RVS, JT, RF and FM contributed to experimental procedures, including embryo production, manipulations, and molecular analysis. RPN performed bioinformatic analysis. LA, RPN, RF, FM, and LS interpreted the bioinformatic results. LA, JT, RPN, RF, and LS wrote the manuscript. All authors contributed to the manuscript revision, and read, and approved the submitted version.

## Acknowledgments

The authors thank Drs. Patrick Blondin and Remi Labrecque from BOVITEQ Inc. for their scientific and technical input. The authors also The authors acknowledge to the server infrastructure of Soroban (SATREPS MACH – JPM/JSA1705) at Centro de Modelación y Computación Científica at Universidad de La Frontera..

## Supplementary Material

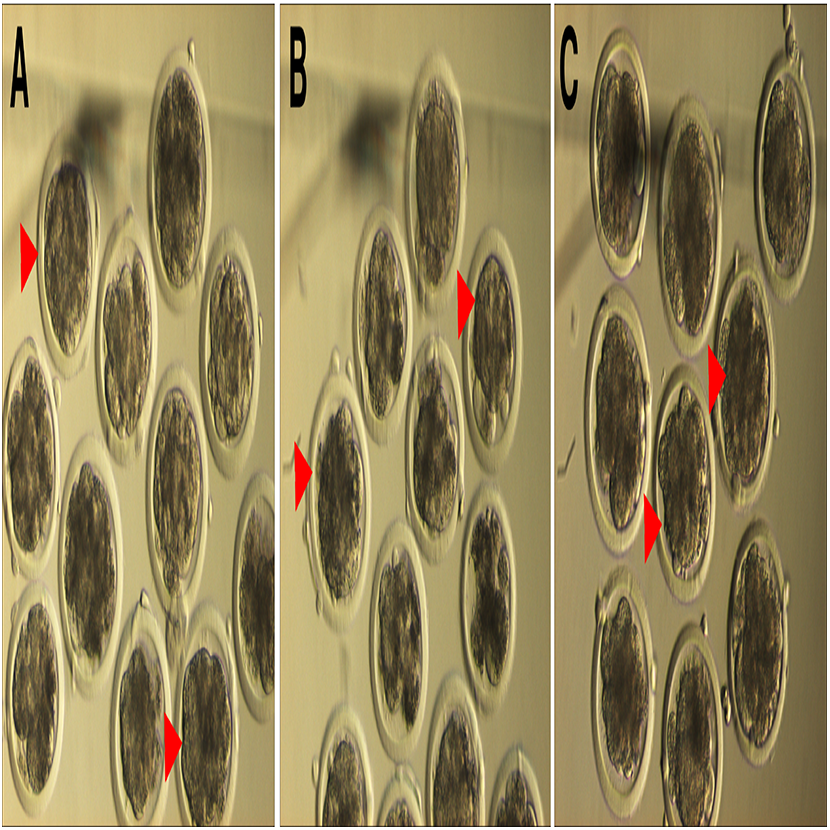

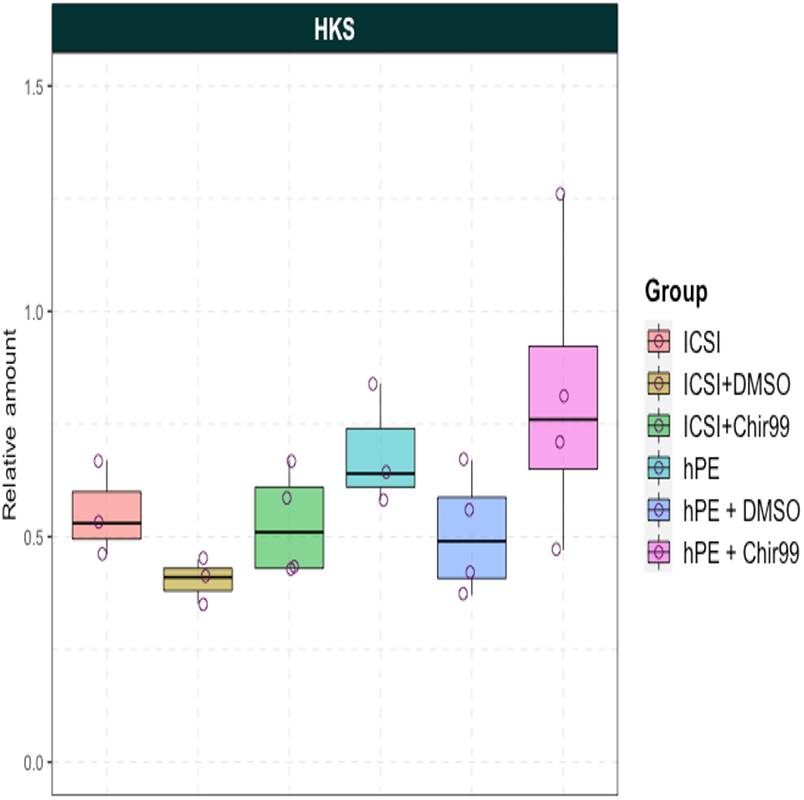

