## Supplemental material for "Haploid androgenetic development in bovines reveals imbalanced WNT signaling and impaired cell fate differentiation"

**Table S1**. Details of primers used for RT-qPCR.

| Details of primers used for RT-qPCR analysis. | | | |
|---|---|---|---|
| **Gene** | | **Primer Sequence (5′−3′)** | **Fragment Size (bp)** |
| ***GAPDH*** | **F:** | TCTGGCAAAGTGGACATCG | 60 |
| | **R:** | GACCATGTAGTGAAGGTCAATGAA | |
| ***SF3A1*** | **F:** | CGAGTTCAAGGAGGGCAAGG | 140 |
| | **R:** | TCTTTGGGCACAATGGTTTCT | |
| ***ACTB*** | **F:** | GCGGACAGGATGCAGAAA | 89 |
| | **R:** | ACGGAGTACTTGCGCTCAG | |
| ***POU5F1*** | **F:** | CGAAAGAGAAAGCGGACGAG | 178 |
| | **R:** | TTGATCGTTTGCCCTTCTGG | |
| ***KFL4*** | **F:** | CCAAAGAGGGGAAGACGGTC | 283 |
| | **R:** | TGTGTAAGGCGAGGTGGTCC | |
| ***SOX2*** | **F:** | CACATGAACGGCTGGAGCAA | 163 |
| | **R:** | AGGTCTGCGAGCTGGTCATAG | |
| ***CDX2*** | **F:** | CAGCCAAGTGAAAACCAGGA | 183 |
| | **R:** | CTTTGCTCTGCGGTTCTGAA | |
| ***CTNNBL1*** | **F:** | TCTGTTCGATCCTCGCTTCC | 179 |
| | **R:** | CATGTCGTGCTTTTCCCCTT | |
| ***YAP*** | **F:** | GCTCAGCACCTTCGACAGTC | 219 |
| | **R:** | CGAGGCCGAATTCATCATGT | |
| ***AXIN2*** | **F:** | GGGAGAAATGCGTGGATACTT | 139 |
| | **R:** | TTGGAGACGATGCTGTTGTTC | |
| ***NANOG*** | **F:** | GCAAACGTCATCTGCTGACAC | 174 |
| | **R:** | TGTTTCTTGACCGGGACCGT | |
| ***GSK3B*** | **F:** | ATTGCACTTTGTAGCCGTCT | 245 |
| | **R:** | GTGGTATTTGAAGGAGCTGA | |

**Table S2.** Morphological assessment of biparental, haploid parthenogenetic and haploid androgenetic embryos at Day-8 (192 h) of culture in presence or not (DMSO) of CHIR99021.

| Group | Blastocyst morphology | | Total |
|---|---|---|---|
| | Early | Expanded/ Hatched | |
| ICSI+DMSO | 39 (63%) | 23 (37%) | 62 (100%) |
| ICSI+Chir99 | 14 (64%) | 8 (37%) | 22 (100%) |
| hPE+DMSO | 24 (67%) | 12 (33%) | 36 (100%) |
| hPE+Chir99 | 18 (58%) | 13 (42%) | 31 (100%) |
| hAE+DMSO | 13 (100%) | 0 (0%) ** | 13 (100%) |
| hAE+Chir99 | 24 (80%) | 6 (20%) | 30 (100%) |

ICSI, intracytoplasmic sperm injection embryos cultured in absence (ICSI + DMSO) or presence of Chir99021 (ICSI + Chir99). hPE, haploid parthenogenetic embryos cultured in absence (hPE + DMSO) or presence of Chir99021 (hPE + Chir99). hAE, haploid androgenetic embryos cultured in absence (hAE+ DMSO) or presence of Chir99021 (hAE + Chir99) (**$p < 0.01$).

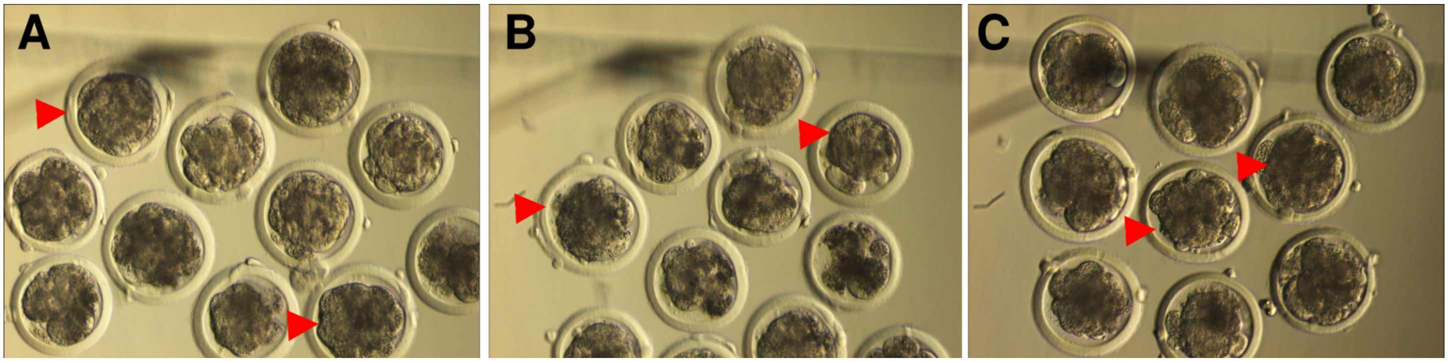


**Fig S1.** Representative images of the morphological quality of day-6 morula stage embryos used for RNA-sequencing. A, intracytoplasmic sperm injection; B, haploid parthenogenetic embryos; C, haploid androgenetic embryos. Red arrowheads indicate selected morphology.

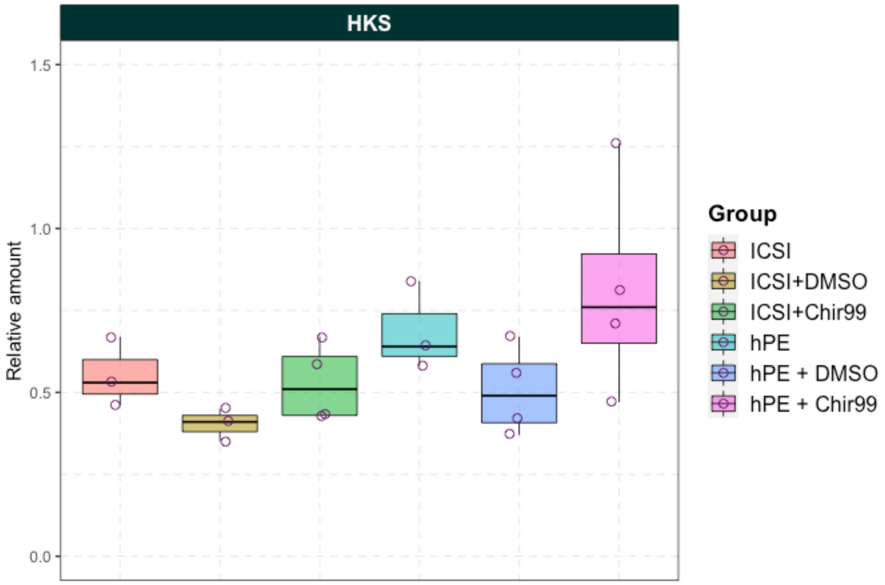


**Fig. S2.** Relative transcript levels of reference genes (HKS, housekeeping) from blastocyst stage embryos (day 8) obtained by ICSI and haploid parthenogenetic activation cultured without (DMSO, 0.01%) or with CHIR99021. ICSI, intracytoplasmic sperm injection embryos; ICSI + DMSO, intracytoplasmic sperm injection embryos cultured in presence of 0.01% DMSO; ICSI + DMSO, ICSI + Chir99, intracytoplasmic sperm injection embryos cultured in presence of Chir99021; hPE, haploid parthenogenetic embryos; hPE + DMSO, haploid parthenogenetic embryos cultured in presence of 0.01% DMSO; hPE +Chir99, haploid parthenogenetic embryos cultured in presence of Chir99021.
